# A fluid-walled microfluidic platform for human neuron microcircuits and directed axotomy

**DOI:** 10.1101/2023.10.14.562004

**Authors:** Federico Nebuloni, Quyen B. Do, Peter R. Cook, Edmond J. Walsh, Richard Wade-Martins

## Abstract

In our brains, different neurons make appropriate connections; however, there remain few *in vitro* models of such circuits. We use an open microfluidic approach to build and study neuronal circuits *in vitro* in ways that fit easily into existing bio-medical workflows. Dumbbell-shaped circuits are built in minutes in standard Petri dishes; the aqueous phase is confined by fluid walls – interfaces between cell-growth medium and an immiscible fluorocarbon, FC40. Conditions are established that ensure post-mitotic neurons derived from human induced pluripotent stem cells (iPSCs) plated in one chamber of a dumbbell remain where deposited. After seeding cortical neurons on one side, axons grow through the connecting conduit to ramify amongst striatal neurons on the other – an arrangement mimicking unidirectional cortico-striatal connectivity. We also develop a moderate-throughput non-contact axotomy assay. Cortical axons in conduits are severed by a media jet; then, brain-derived neurotrophic factor and striatal neurons in distal chambers promote axon regeneration. As additional conduits and chambers are easily added, this opens up the possibility of mimicking complex neuronal networks, and screening drugs for their effects on connectivity.

## Introduction

Various *in vitro* methods have demonstrated the requirement for complex culture systems to support neuronal maturation and the manifestation of associated disease ^1–3^. Microfluidic approaches have yielded particularly promising results, as they enable isolation of different cellular compartments (e.g., somas, dendrites, axons) ^4,5^ . Compared to conventional *in vitro* cultures, they also permit precise control of cellular environments, and have proven useful in studies on neurotoxicity ^6^ and electrical connectivity ^7,8^. Nevertheless, conventional microfluidic devices have limitations that are often attributed to the materials used for fabrication ^9^; they are usually made of a plastic elastomer (polydimethylsiloxane, PDMS) firmly bonded to a glass substrate, and cells are buried in chambers bounded by solid walls that prevent insertion of the standard experimental tools used by neurobiologists (e.g., cell scrapers, patch-clamping pipettes). Consequently, neurobiologists must employ alternative protocols to use them that are different from their familiar ones.

Recently, Walsh et al. (2017) introduced ‘fluid-walled microfluidics’; this overcomes some of these limitations by removing most solid boundaries ^10^. It is a form of open microfluidics ^11^ that exploits properties of fluids at the microscale to confine aqueous environments using interfaces (i.e., fluid walls) between immiscible liquids (in this case, cell growth medium and a bio-inert fluorocarbon, FC40). Circuits can be built in minutes in standard Petri dishes, and – unlike solid walls that cannot be pierced by pipets – fluid walls allow direct access to cells everywhere in circuits. These walls re-heal automatically when pipets are withdrawn, they can be destroyed and/or reshaped without damaging cells within them ^12,13^, and are so transparent that cell morphology can be monitored using standard microscopes ^14^. These features motivate the use of this technology here.

The cortex and striatum, together with the basal ganglia and thalamus, play an important role in regulating voluntary movement, learning, executive function, and emotion ^15^. Cortical neurons (CNs) project axons toward the striatum where medium spiny neurons (MSNs) constitute up to 95% of striatal subtypes. Connectivity between CNs and MSNs is directional and monosynaptic, whilst MSNs communicate with CNs indirectly via downstream circuits, particularly the basal ganglia ^16^. This oriented arrangement of cortico-striatal projections is critical for their functioning but is often oversimplified by *in vitro* co-cultures.

Studies of axonal outgrowth, axotomy, and subsequent regeneration have been facilitated by the use of compartmentalised chambers that allow separation of cell bodies from their axons ^17,18^; such platforms can also allow pharmacological screening of factors acting specifically on distal axons ^17,19^. In this study, we exploit the advantages of fluid walls to establish a proof-of-concept model that recreates appropriate arrangements of human CNs and MSNs. We then go on to develop a method for targeted and localised axotomy of CNs, and compare the effects of pro-regenerative conditions on axonal regrowth ^19–21;^ brain-derived neurotrophic factor (BDNF) and postsynaptic MSNs both have positive effects. In combination, these approaches provide a method for screening drugs promoting developmental outgrowth of axons and regeneration of damaged ones.

## Results

### Jet-printing of fluid-walled micro-circuits shaped like dumbbells

The fluid-walled environment is created in standard polystyrene Petri dishes (6 cm) by ‘jet-printing’ (A) ^22^. A thin layer of cell-culture medium is overlaid by an immiscible, bio-inert, and clear fluorocarbon, FC40 (Figure 1Ai). Next, a submerged jet of FC40 (Figure 1Aii) sweeps medium off the plastic substrate to leave FC40 locally pinned to the dish (Figure 1Aiii). The jetting nozzle (held by a 3-way traverse) now moves laterally above the substrate to draw the outline of the desired pattern which is held to the dish by interfacial forces acting between the two immiscible phases and the substrate. Here, we fabricated a 7×3 array of dumbbell-shaped circuits in each dish; each dumbbell comprises two square chambers (3 mm) connected by a thin conduit (∼0.2 mm wide, 1 mm long, <10 µm high; Figure 1B).

**Figure 1.**
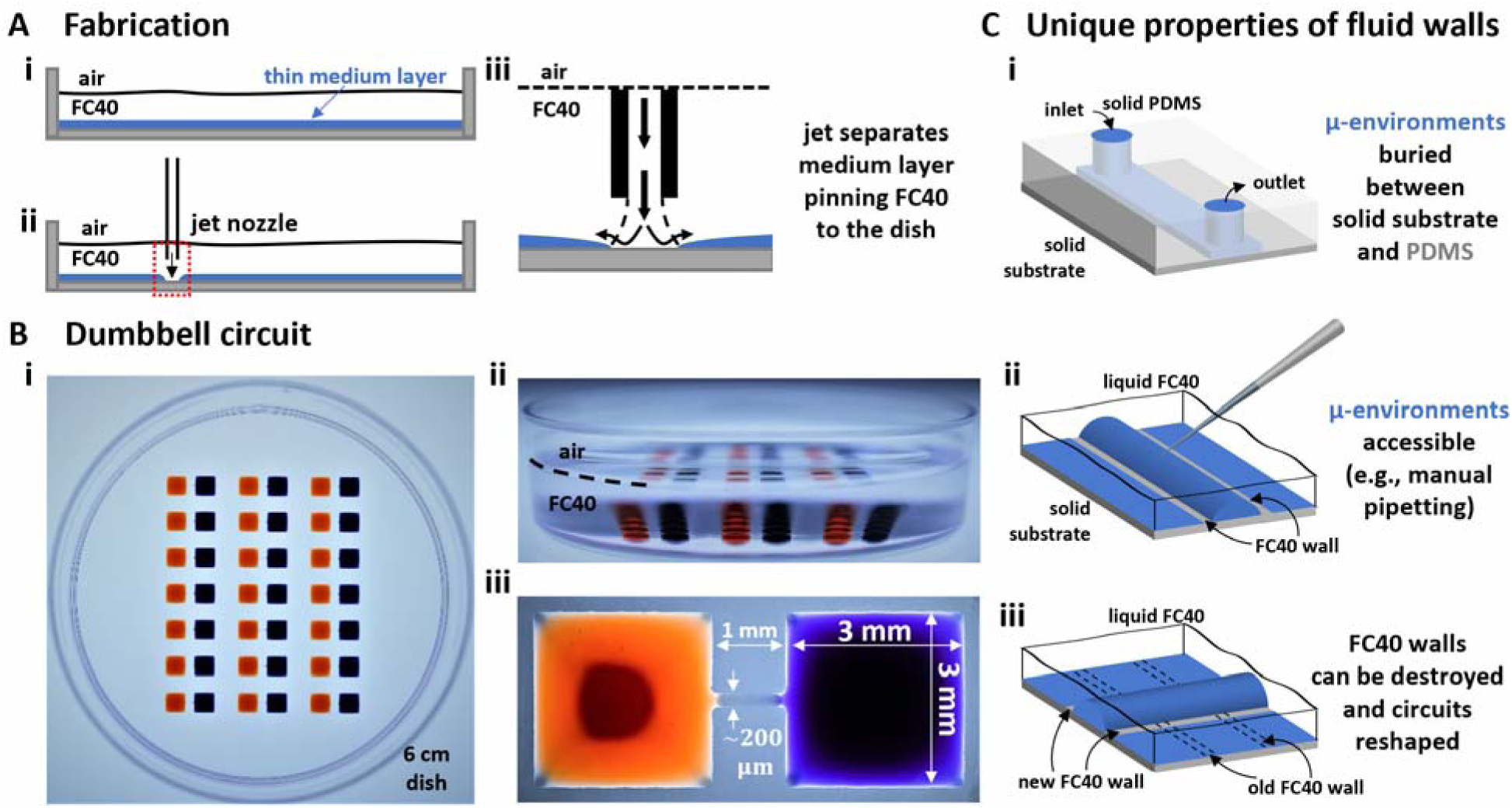
Fabrication and operation of fluid-walled circuits. **(A)** Fabrication. (i) In a standard polystyrene Petri dish, a thin layer of cell-culture medium is overlaid b an immiscible, transparent, and bio-inert fluorocarbon (FC40). (ii) Additional FC40 is jetted (480 μl/min) through a nozzle mounted on a 3D-traverse. (iii) The submerged jet sweeps away medium, to leave fluid walls of FC40 pinned to the dish along the path of the traverse. **(B)** Array of dumbbells after filling each with red and blue dyes. (i,ii) Top and side views of dish. (iii) Zoomed-in image of one dumbbell (conduit length = 1 mm). **(C)** Comparing properties of solid PDMS walls used in conventional devices, with fluid ones. (i) Access to a conventional device is only through inlet and outlet ports. (ii) Medium can be pipetted into or out of any point in a fluid-walled circuit as liquid-liquid interfaces are easily pierced to re-heal automatically on withdrawal. (iii) Fluid walls can be destroyed at any time, and different ones recreated on demand.

Compared to conduits in conventional devices with solid walls – where access is limited to inputs and outputs (Figure 1Ci) – all parts of dumbbells are accessible through fluid ceilings (Figure Cii). Media and/or cells were added to, and removed from, chambers by lowering a dispensing needle (also held by the traverse and connected to a syringe pump) through the FC40 until its tip was near the surface of the medium (i.e., 200 µm above the bottom of the dish). Existing walls/ceilings can also be destroyed and rebuilt (Figure 1Ciii), so circuits can be reconfigured during experiments ^12^. Additionally, fluid walls are freely permeable to vital gases, so cells in these circuits are grown in conventional CO_2_ incubators. Finally, the refractive index of FC40 (1.29) almost matches that of water (1.33), and this permits undistorted imaging with standard microscopes ^14^.

### Local pressures in dumbbells

In all experiments, when cells are deposited in chambers, we require they remain there. This is impossible to achieve by rapid deposition into a newly-fabricated dumbbell, as this induces flow that carries them into the conduit (and perhaps into the other chamber). Therefore, we begin by describing how local pressures within dumbbells can be manipulated.

First consider a 1 µl drop sitting in a dish filled with FC40; the drop is shaped like the cap of a sphere, as interfacial forces minimise the contact area of medium with the immiscible fluorocarbon. Then, the pressure (P) at the base of the drop is defined by the Laplace pressure across the medium:FC40 interface, plus the hydrostatic head of overlying medium and FC40. The Young-Laplace equation gives Laplace pressure, *LP* = *y*(^1^1_*R*1_ + ^1^1_*R2*_), where *γ* is the interfacial tension, and *R*_1_ and *R*_2_ are two orthogonal radii describing the curvature of the liquid wall/ceiling. Assuming our chambers have circular footprints like the drop (so *R*_1_ = *R*_2_ =*R* = ^*u*^^2^ ^+*h2*^ 1_*2h*_, with *a* being the radius of the chamber footprint),

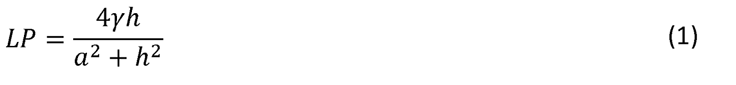

The combined pressure, *P*, at the base of a chamber then includes the two hydrostatic heads, so

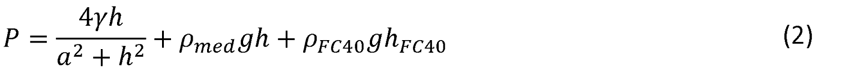

where *µ*_*med*_ and *µ*_*FC40*_ indicate density of medium and FC40 respectively, !J is gravitational acceleration, *h* is drop height, and *h_FC40_* the height of the overlay. However, when analysing the pressure difference between our two chambers (ΔP), the contribution of *h_FC40_* can be expressed as the height difference of the two chambers. Therefore:

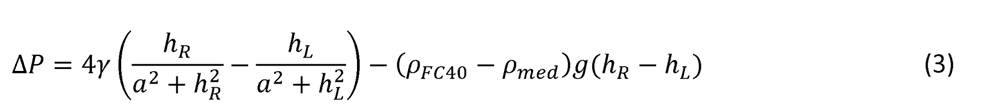

where *h_R_* and *h_L_* indicate right and left chamber height, respectively. *h_R_* and *h_L_* represent the only variables in Eq. (3) as they depend on the volume (V) infused into each chamber, and

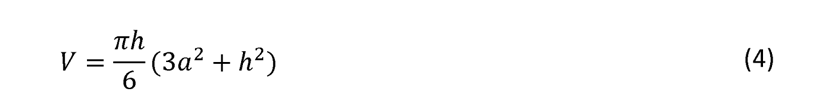

This means the pressure difference can be controlled simply by controlling chamber volume.

## Ensuring cells remain where deposited

After fabrication, chamber volumes are minimal, walls/ceilings are almost flat (so Laplace pressures are almost negligible), local pressures throughout a dumbbell are roughly equal, and the system is in equilibrium (Figure 2Ai, 2Bi). Adding 4 *µ*l into the right-hand chamber followed by 1 μl into the left-hand one generates a pressure difference that induces leftward flow through the conduit (Figure 2Aii, 2Bii). As time passes, the system equilibrates and volumes equalise (Figure 2Aiii, 2Biii). Similarly, after adding 4 *µ*l blue dye into a right-hand chamber and 1 *µ*l red dye into the left-hand one (Figure 2Ci,ii), red dye is confined to the left-hand-side for at least 24 h (Figure 2Ciii; note the conduit is filled with blue dye, and the left-hand chamber contains both dyes). Quantification of chamber pressures and volumes over time confirm that both converge towards equilibrium values, and that blue pressure is always greater than red pressure over 24 h (Figure 2D). This confirms that if the right-hand chamber is pre-filled with 4 *µ*l before adding 1 *µ*l to the left-hand one, there cannot ever be flow rightward. We used this approach to ensure that when cells are seeded in a selected chamber, they remain there.

**Figure 2.**
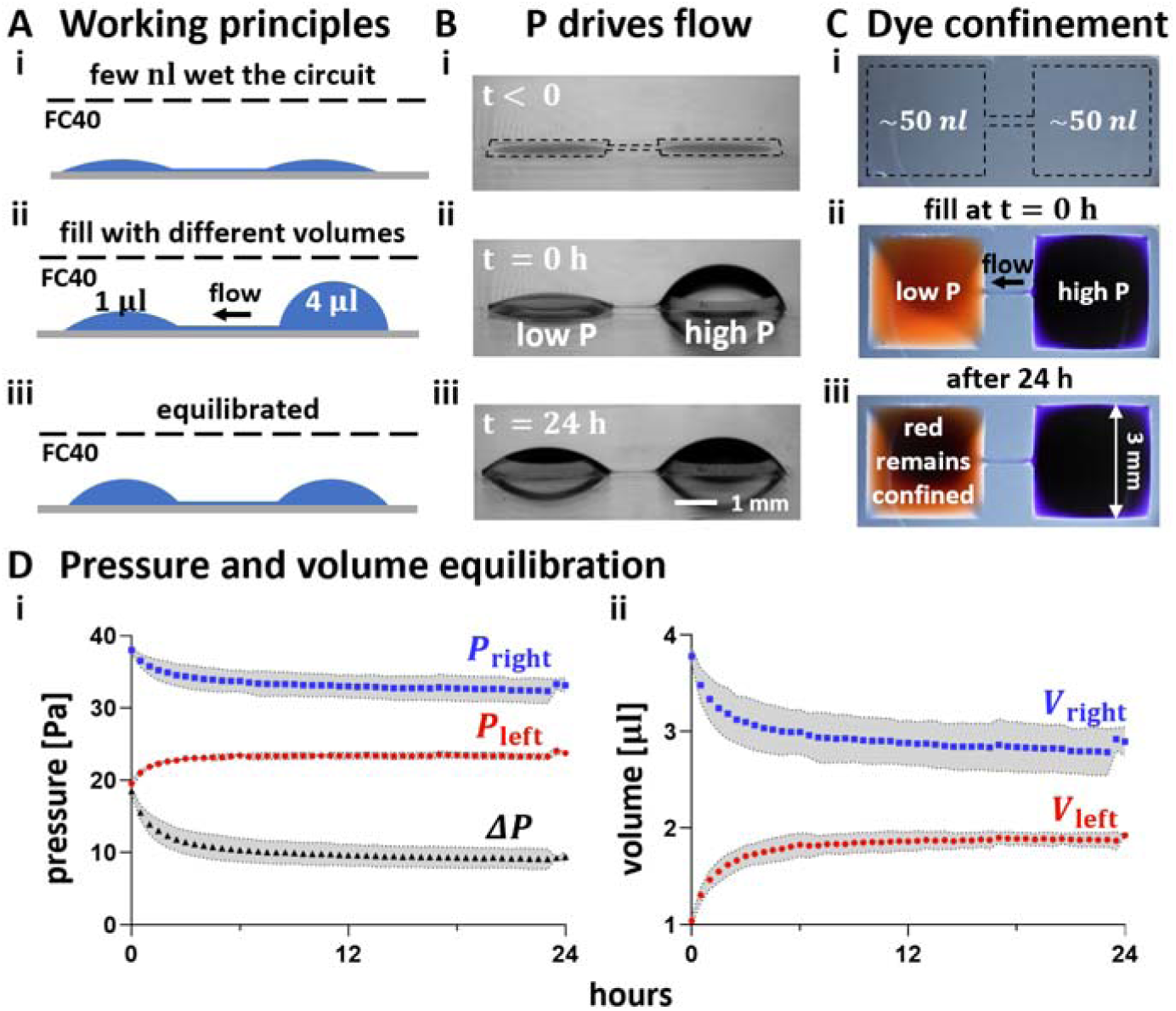
Ensuring liquid pipetted into the left of a dumbbell remains there. P = pressure. **(A)** Principles. (i) A dumbbell is flow-free after fabrication. (ii) Medium (4 *µ*l) is added to the right-hand chamber; this generates a high local Laplace pressure. When 1 *µ*l is immediately added to the left-hand chamber (creating a lower local Laplace pressure), resulting left-ward flow through the conduit prevents any of the 1 *µ*l from moving to the right. (iii) Eventually the system equilibrates, and flow ceases. **(B)** Pressure difference drives flow. (i) Before filling. (ii) Immediately after filling with different volumes. (iii) After 24 h, chamber volumes have almost equalised due to leftward flow. **(C)** After adding 4 *µ*l blue dye to the right and 1 *µ*l red dye to the left, no red dye flows rightward (top views). (i) Before adding dyes (dotted line marks dumbbell footprint). (ii) Immediately after adding dyes. (iii) After 24 h, red dye remains confined in its chamber (which now also contains blue dye). **(D)** Changes in pressure (i) and volume (ii) of right-(blue) and left-hand (red) chambers determined after measuring chamber heights and calculating values using Eq. 1 (and) and Eq. 3, respectively. Black curve: pressure difference between chambers. Each dot represents the mean value of 3 technical replicates, and grey areas the associated standard deviations.

When both chambers have equal volumes, their internal pressures are also equal – and so there is no flow in either direction. As a result, mass transport between chambers only occurs by diffusion to generate a concentration gradient in the conduit, with the steepness and duration of the gradient depending on dumbbell geometry and diffusion constant (see Supplementary Information). Supplementary Figure 1 and Supplementary Table 1 show how such a gradient of BDNF changes over time.

## Axons outgrow from CNs through the conduit to the distal chamber

Axon pathfinding is led by a variety of molecular cues, both intrinsic ^23^ and target-derived ^24^. While the roles of intrinsic factors have been investigated using dissociated cultures of neurons *in vitro* ^21,25^, less is known about target-derived signals due to difficulties in recreating the required micro-environments around neurons. Here, we developed a microfluidic model that recreates such an environment; we exploited the intrinsic physics of fluid-walled dumbbells (Figure 2), and established a workflow (Supplementary Figure 2) that confines CNs in the left-hand chamber as axons grow through the conduit to the right-hand (distal) one.

Figure 3A provides an overview of the 3 major workflows that are now used. In one (yellow arrow), human induced pluripotent stem cells (iPSCs) were induced to differentiate into (post-mitotic) CNs using proven methods in standard well plates ^26^. In a second (red arrow), human iPSCs were similarly induced to develop into post-mitotic MSNs, again using conventional methods ^27^. In a third (blue arrow), fluid-walled dumbbells were jet-printed in 6 cm polystyrene Petri dishes, then CNs and MSNs plated into dumbbells.

**Figure 3.**
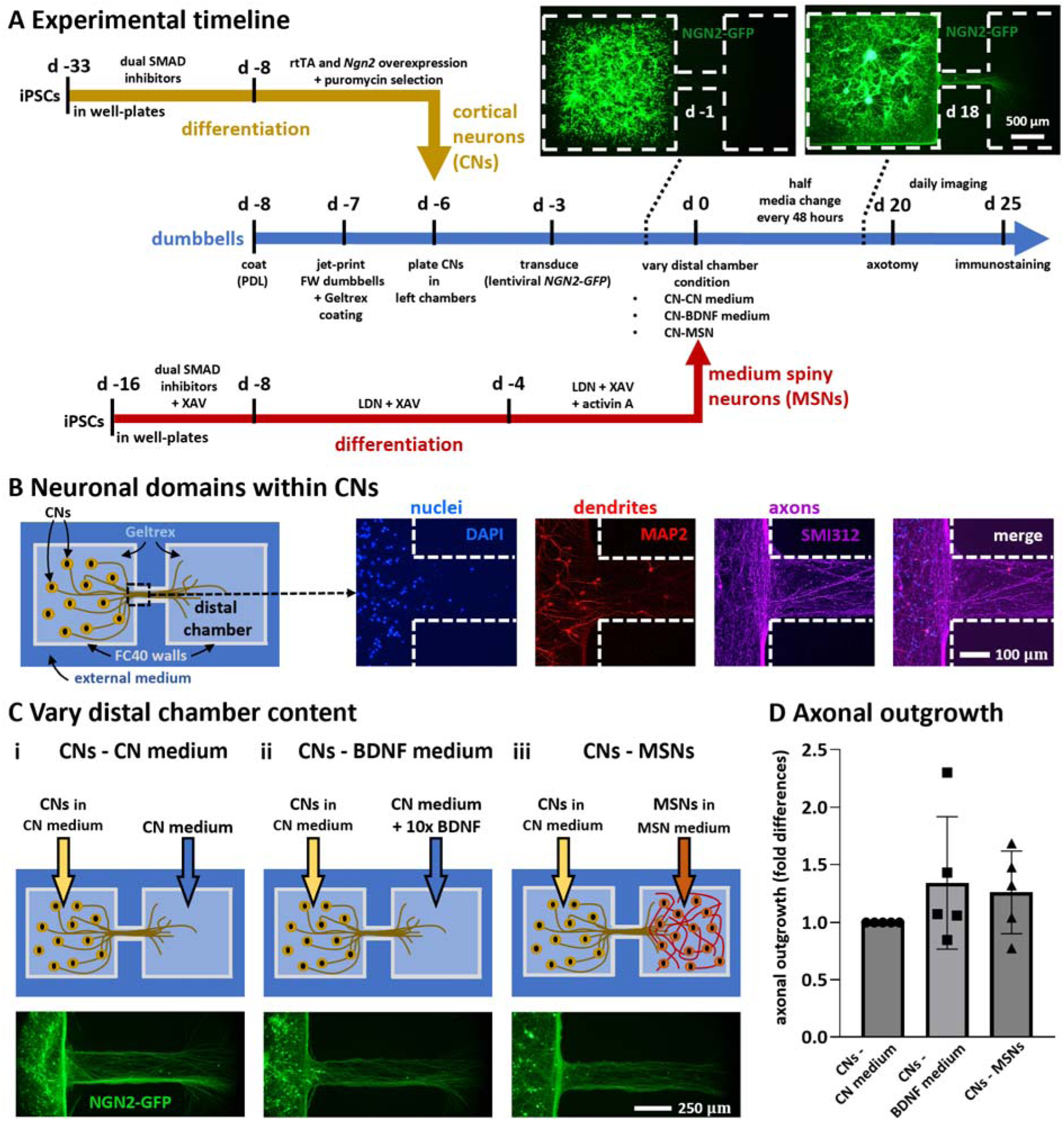
Timelines and culture conditions used to model and study unidirectional outgrowth of cortical neurons. **(A)** Timelines. Yellow and red lines show protocols for generating CNs and striatal MSNs from iPSCs using standard methods; blue line describes those involved in making and operating fluid-walled dumbbells. Insets: after plating transduced CNs in left-hand chambers, live-cell images (collected on d -1 and 18) show NGN2-GFP fluorescence in parts of dumbbells (dotted white lines show circuit edges). **(B)** Immunostaining shows compartmentalisation of neuronal domains in CNs (d 25). Cartoon: CNs in left-hand chamber, CN medium (but no MSNs) in distal one. Nuclei (DAPI) and MAP2-positive dendrites are confined within the chamber, while only SMI312-positive axons grow into the conduit (dotted white lines show circuit edges). **(C)** Effects of varying distal-chamber content on axonal outgrowth. In all cases, CNs expressing NGN2-GFP are plated in CN medium in left-hand chambers. Top: cartoons indicating chamber contents. Bottom: live-cell epifluorescence images of left end of conduit (d 20); axons extend into conduits in all three cases. (i) Monoculture control. (ii) Positive control with regenerative medium in distal chamber (CN medium + 10-fold higher concentration of 100 ng/ml BDNF. (iii) MSNs + MSN medium in distal chamber (this condition attempts to promote connectivity between cortex and striatum seen in vivo). **(D)** Quantitative analysis of axonal outgrowth seen in (C). Outgrowth (fold difference) is difference in area covered by GFP-expressing neurites in conduits between d 0 and d 20, normalised for GFP-positive conduit area on d 0 as a function of the number of cells, and expressed relative to their control. Each dot represents a healthy control-derived line from one differentiation. N = 2 iPSC lines, n = 2-3 differentiations/line. One-way ANOVA with Bonferroni correction; p > 0.05.

In all experiments that will be described, CNs are deposited into left-hand chambers of dumbbells (using conditions established in Figure 2), and transduced 3 days later with lentiviruses encoding (tetracyclin-inducible) neurogenin-2-GFP (NGN2-GFP) to allow live-cell visualisation of transduced CNs. During subsequent culture (with CN medium in the right-hand chamber but no cells), axons (now expressing NGN2-GFP) grow through the conduit into the distal chamber (Figure 3A, insets; compare live-cell images on days -1 and 18). In some cases, different media and/or MSNs were deposited on d 0 in the right-hand (distal) chamber. To ensure that deposited MSNs remained where deposited, we again exploited the approach described in Figure 2, except that now it is the left-hand chamber that was pre-filled with 4 *µ*l before 1 *µ*l of cell suspension is plated in the distal one.

As we wish to replicate the spatial organisation of CNs and MSNs in vivo, we require that CN somas remain confined to the left-hand chamber, and that only axons grow into the conduit. We confirmed successful compartmentalisation by immunostaining (d 25) using domain-specific markers: DAPI-labelled nuclei and MAP2-positive dendrites were confined to the left-hand chamber, while the conduit was populated with SMI312-positive axons (Figure 3B).

We have seen that after plating in the left-hand chamber, post-mitotic CNs – with standard CN (maturation) medium in both chambers – started projecting axons into the conduit between d -6 to d 0. We then varied contents of the distal chamber at d 0 to see what effects concentration gradients of diffusing molecules along the conduit have on axonal outgrowth. This was tested using three different conditions. First, a control where CNs project axons towards distal CN medium – so there is no chemical gradient in the conduit (CNs-CN medium condition). A second condition where the distal chamber was filled with CN medium supplemented with 10x concentration (100 ng/ml) of BDNF (CNs-BDNF medium condition). Third, a more physiological condition where CNs extended their projections towards a population of MSNs (CNs-MSNs condition).

Axonal outgrowth in each of the 3 conditions was quantified as the difference of areas covered by GFP-expressing neurites in conduits measured at d 0 and d 20 (ΔA = A_d20_ -A_d0_). As ΔA may vary depending on the number of cells in each chamber making comparison between dumbbells difficult, we divided it by A_d0_ to normalise results decoupling them from the effective count of CNs seeded. Additionally, to compare results from different batches of differentiated iPSCs, axonal outgrowths in the CNs-BDNF medium and CNs-MSNs conditions were expressed as the fold-difference of the CNs-CN medium control of the respective batch. Perhaps surprisingly, neither condition significantly increased axons outgrowth (Figure 3D; p > 0.05, one-way ANOVA with Bonferroni correction) suggesting it occurred independently of exogenously-supplied BDNF or MSN-derived molecules.

## A unidirectional circuit between cortex and striatum

The CNs – MSNs condition described above provides an *in vitro* model for possible connectivity between cortex and striatum. This prompted a more detailed immunolabelling analysis on d 25 (Figure 4). Note here that NGN2-GFP (now detected using an anti-GFP antibody) is a CN-specific marker, and DARPP32 is a MSN-specific one; MAP2 and SMI312 mark respectively dendrites and axons in both cell types. As expected, CNs in the left-hand chamber expressed NGN2-GFP plus MAP2, but no (striatal-specific) DARPP32 (Figure 4A,B). The right-hand end of the conduit contained no nuclei stained with DAPI (confirming somas remain where plated), but many SMI312-positive axons that appeared to invade the distal chamber with its MAP2-positive dendrites (Figure 4C). The distal chamber was populated by many (CN-derived) NGN2-GFP-positive axons that ramified amongst MSNs expressing (striatal-specific) DARPP32 plus MAP2. These results are consistent with no spill-over of somas from either chamber, and unidirectional growth of CN axons into the distal chamber where they ramify amongst MSNs.

**Figure 4.**
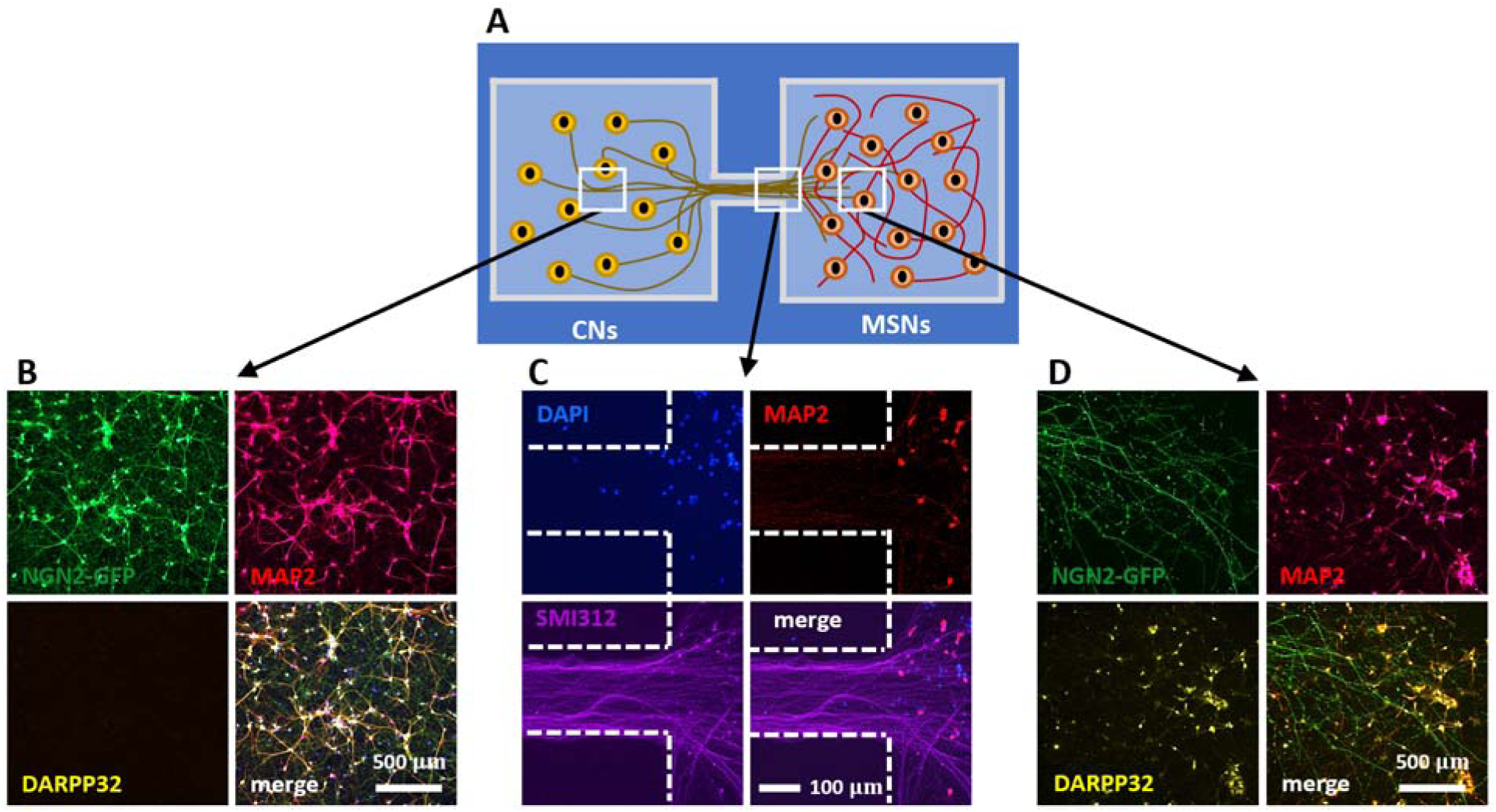
Modelling of unidirectional pathway between cortex and striatum. Cortical axons marked with NGN2-GFP grow from the left chamber through the conduit into the distal chamber containing MSNs. (DAPI – blue – nuclei, MAP2 – red – dendrites, SMI312 – purple – axons, NGN2-GFP – green – transduced CNs, DARPP32 – yellow – MSN marker; merges indicated). **(A)** Top-view schematic of the cortex-striatum circuit in a fluid-walled dumbbell. CNs are seeded in the left chamber and cultured for 6 days, then MSNs are plated in the distal (right) chamber; after growth for another 25 days, cells are immunolabelled. White boxes indicate areas imaged below using various markers. **(B)** MAP2-positive dendrites and NGN2-GFP are found throughout the CN chamber (but not DARPP32-expressing MSNs). **(C)** MSNs nuclei remain where seeded and develop MAP2-positive dendrites; they are joined by SMI312-positive CN axons from the left-hand chamber. **(D)** Axons containing NGN2-GFP from transduced CNs are intertwined amongst MAP2-and DARPP32-positive neurons in the striatal chamber.

## Axotomy using a micro-jet

We next used a micro-jet to sever axons projecting from CNs through a conduit into an empty distal chamber. The process involved three steps. First, fluid walls were destroyed by removing (manually by pipet) the FC40 overlay, and adding ∼5 ml medium to the dish; CNs and axons remained attached to the substrate (Figure 5Ai). Second, a submerged jet of medium (emitted from the same nozzle used for FC40 jet-printing) was moved perpendicularly by the traverse across the axons to sever them (Figure 5Aii). Third, new fluid walls were built just outside the original ones to recreate a slightly-larger dumbbell (Figure 5Aiii). To do so, most medium was gently removed to leave a thin layer, fresh FC40 was overlaid, and a new dumbbell jet-printed around the original one so that the FC40 stream did not impinge on existing attached cells or axons (Supplementary Figure 2C). This technique allows targeted axotomy of a chosen segment within an axon (Figure 5Aiv). Moderate-throughput axotomy in each of the 21 original dumbbells in one dish is achieved in <90 sec (Figure 5Av). Comparison of live-cell images of cortical axons expressing NGN2-GFP taken before and after axotomy revealed how local damage was, with the jet clearing the axotomized area of most cellular material (Figure 5Bi,ii).

**Figure 5.**
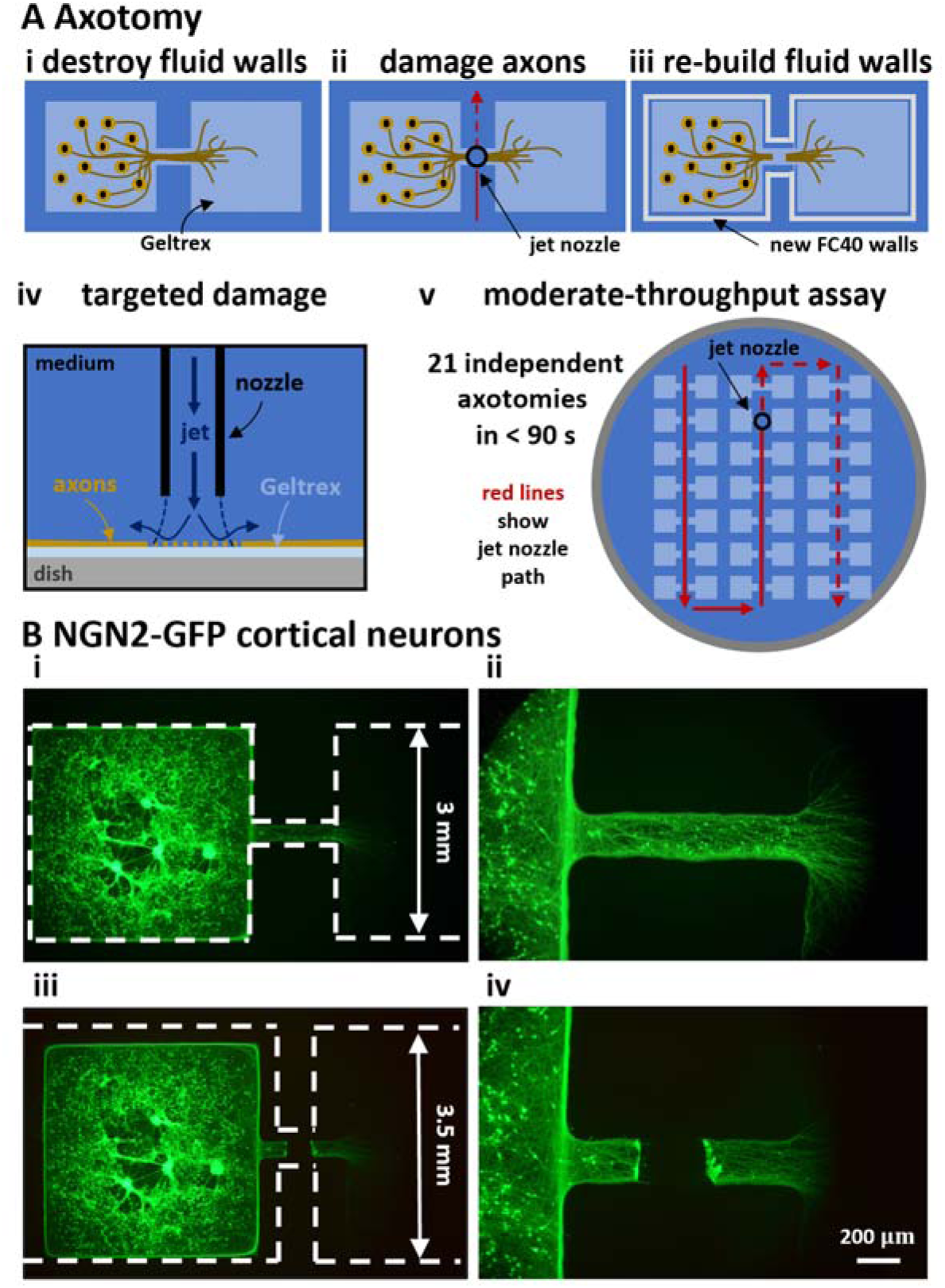
Axotomy assay on fluid-walled microfluidics. **(A)** Schematics illustrating localized axotomy. (i) Fluid walls are destroyed; cells remain attached to Geltrex within the dumbbell footprint. (ii) The traverse moves a nozzle that jets medium onto axons and cuts them. (iii) New fluid walls are jet-printed (using an FC40 jet) to form a new dumbbell slightly larger than the original one; after filling the new dumbbell, axons regrow. (iv) Higher-magnification schemati of the submerged media jet in action targeting axons to generate localised damage. (v) Overview. Red lines illustrate the path followed by the medium jet nozzle while damaging 21 dumbbells (7×3 array) in a single 6 cm Petri dish in less than 90 seconds. Solid lines indicate the path already covered; dashed lines show future nozzle positions. **(B)** Representative live-cell fluorescent images of CNs expressing NGN2-GFP pre-and post-axotomy. (i,ii) Dashed lines mark edges of dumbbell footprints before and after axotomy. (iii, iv) Higher magnification images taken before and after axotomy.

## Regeneration after axotomy

After printing new dumbbells around damaged cultures, the three conditions were re-established with equal volumes of correct media in each respective chamber (Figure 6Ai), and then regeneration of axons was monitored for 5 more days (till d 25). Representative immunostaining on d 25 shows the axotomized area of the conduit refilled with axons only expressing NGN2-GFP and SMI312, but not the dendrite-specific MAP2 (Supplementary Figure 3). As regrowth after axonal damage may involve target-derived signalling ^28,29^, we compared effects of our three conditions on this regeneration (Figure 6Ai).

**Figure 6:**
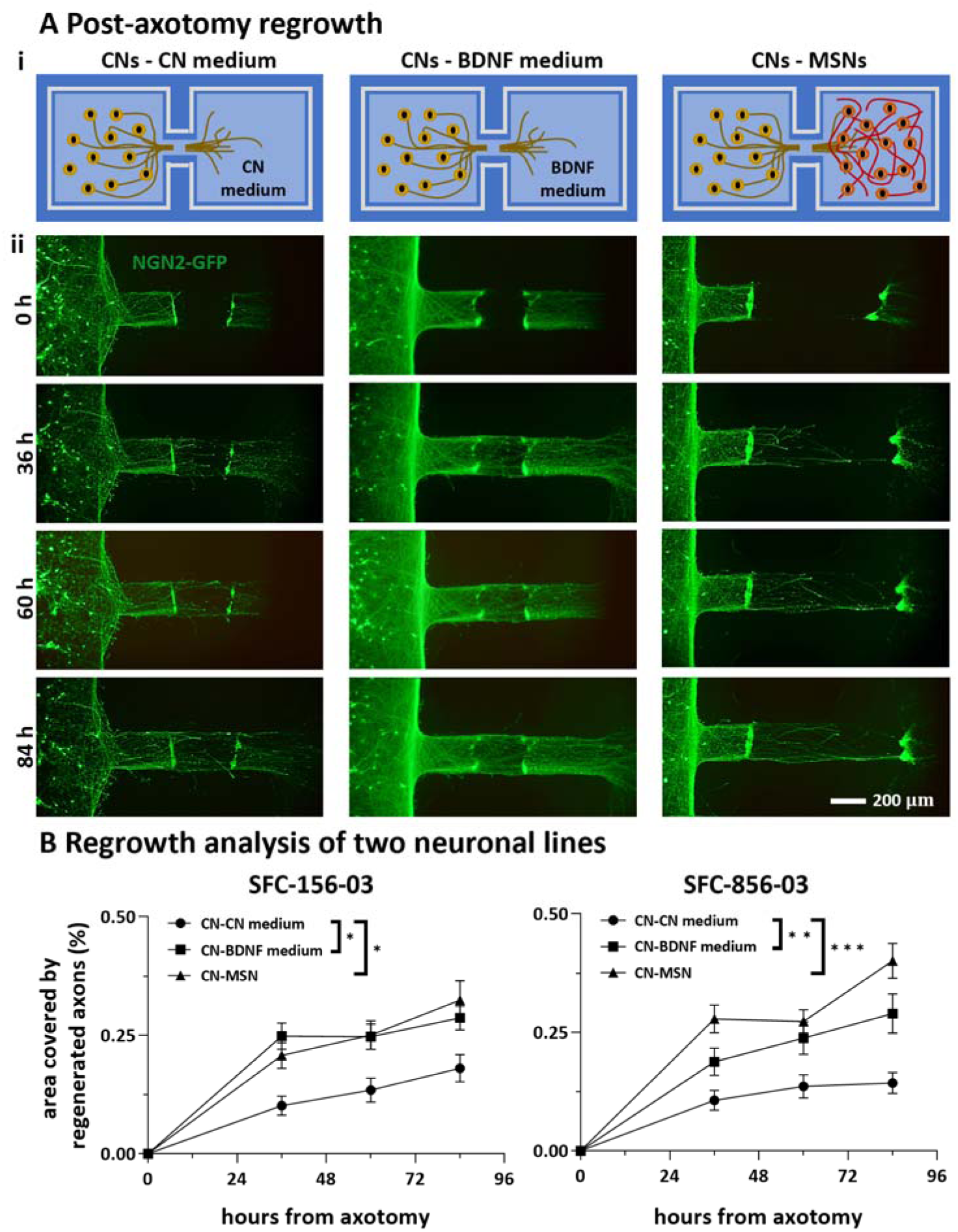
Effects of distant target-derived factors on regrowth of cortical axons after damage. **(A)** Schematic and representative live-cell fluorescent images of CNs expressing NGN2-GFP taken 0-84 h after axotomy. **(B)** Fraction of axotomized area re-covered by axons derived from two different healthy control iPSC lines (SFC-156-003, SFC-856-03). N =2-3 differentiations per iPSC line, * p <0.05, ** p < 0.01, *** p < 0.001, two-way ANOVA with Bonferroni correction test. (SFC156-03: CN-CN medium versus CN-BDNF medium, p > 0.05 at 36 and 60 hours, p < 0.01 at 84 hours; CN-CN medium versus CN-MSN, p <0.0001 at 36 hours, p < 0.001 at 60 and 84 hours; SFC856-03: CN-CN medium versus CN-BDNF medium, p <0.001 at 36 hours, p < 0.05 at 60 and 84 hours; CN-CN medium versus CN-MSN, p < 0.01 at 36 and 84 hours, p < 0.05 at 60 hours; two-way ANOVA with Bonferroni correction test). Furthermore, BDNF and MSNs exert comparable positive effect on cortical axonal regeneration (for both iPSC lines, CN-BDNF medium versus CN-MSN p > 0.05, two-way ANOVA with Bonferroni correction test).

As for outgrowth, regeneration was quantified as the area covered by GFP-expressing neurites growing into a rectangular zone within the cleared area (Figure 6Aii; axon tracts that are incompletely severed or fully cleared are excluded from this analysis). This zone abuts the proximal damage line and has the same width of the original conduit (i.e., 200 μm) while extending 300 μm into the cleared area. Results were normalised against values seen prior to axotomy on d 20 to account for differences in axonal numbers and transduction efficiency. Results obtained with our two different iPSC lines from healthy donors (again with 2-3 differentiations per line) show that both BDNF and MSNs enhance regeneration to roughly the same degree compared to the control (Figure 6B). These results show that severed CN axons can regenerate, and that BDNF and MSNs enhance this.

## Discussion

Our strategic goal is to develop microfluidic methods facilitating the production of, and experimentation on, neuronal circuits of any type *in vitro*; critically, we require that these methods fit easily into bio-medical workflows. We described proof-of-principle experiments illustrating individual steps towards this end. We began by fabricating in minutes dumbbell-shaped micro-circuits in standard Petri dishes (6 cm; Figure 1), and developed conditions ensuring that cells plated in one or other chamber remain where deposited (Figure 2). Next, post-mitotic CNs derived from human iPSCs were seeded in left-hand chambers of the dumbbell, so axons could grow through connecting conduits of 1 mm to distal chambers (Figure 3A,B). Outgrowth of cortical axons was compared against three different targets in the distal chamber (Figure 3C,D); results showed such outgrowth was not enhanced by BDNF. This result aligns with a previous study on mouse primary dorsal route ganglia (DRG) cultures ^30^ in which high levels of BDNF did not boost normal distal growth. Another study ^31^ observed that bathing mouse primary cortical neurons in 50 ng/ml BDNF increased axonal outgrowth by ∼25%. In our study, BDNF was applied distally and the CNs bodies were exposed to a maximum of 15 ng/ml BDNF, allowing for diffusion of BDNF from the distal chamber (see Supplementary Information), likely explaining the difference in results. Although the BDNF signalling pathway is likely conserved across species ^29^, as far as we are aware, this is the first report on the influence of external diffusible factors applied distally on axonal outgrowth of human CNs *in vitro*; hence, modulation by BDNF as well as other neurotrophins will require further investigation. Interestingly, as MSNs also showed no effect on normal healthy outgrowth, we hypothesise axonal outgrowth of post-mitotic human iPSC-derived CNs to be independent of endogenous postsynaptic targets.

We then developed a circuit in which axons project from CNs in the left-hand chamber, through the conduit, and on to ramify amongst MSNs in the distal one (Figure 4). We anticipate such circuits will prove especially useful for studying cortical-striatal connectivity because they are so accessible to biologists. For example, an obvious next step is to examine electrical connectivity using patch clamping ^27^ and/or super-resolution of pro calcium imaging ^32^; our circuits can be built on glass surfaces and incorporated into both workflows without modification.

Similar to previous work ^30,33^, we also developed a moderate-throughput axotomy assay in which axons growing through conduits are severed by a fluid jet (Figure 5). In contrast to existing methods that use mechanical stresses ^34,35^, vacuum aspiration ^19,36,37^ or toxins ^38^ for axotomy - all procedures that might be expected to yield poor reproducibility ^39^ - we hope our (arguably non-contact) method will yield more consistent results. In our assay the quality of the axotomy depends on cells density and age as these parameters affect the thickness of the axonal bundle; ‘cut’ parameters like jet flow rate, nozzle height above the dish, and traverse speed were finely tuned to achieve the desired result. Using this assay, we showed that BDNF or MSNs in distal chambers promote axon regrowth (Figure 6).

Here, the heightened demand for BDNF following axon injury ^40^ might explain the observed positive effect on axonal regeneration despite the neutral effect shown on axonal outgrowth. Co-culture of CNs with their striatal target MSNs also promoted cortical axonal regeneration post-axotomy comparable to that induced by exogenous BDNF, suggesting that the distant postsynaptic target could release pro-regenerative factors acting to facilitate reinnervation.

In conclusion, we have recreated a basic micro-circuit containing human neurons that mimics the unidirectional connectivity seen between cortex and striatum in vivo, as well as developing a moderate-throughput axotomy assay. Our approaches benefit from the intrinsic advantages provided by fluid walls that include ease of circuit fabrication and operation, plus compatibility with existing bio-medical workflows. As additional conduits and chambers can be easily added, we anticipate these approaches will expand the experimental toolkit available for the study of human neuronal networks in health and disease.

## Materials and methods

### Fluorocarbon 40 (FC40)

FC40 was purchased from 3M. FC40^STAR^ (iotaSciences Ltd, Oxfordshire, UK) is a compound treated with a proprietary method to improve formation of fluid walls. Throughout the article, the term ‘FC40’ is used to refer to FC40^STAR^.

### The fluid printer

The printer (iotaSciences Ltd, Oxfordshire, UK) consists of 3D traverse with two printing heads (two blunt needles of different internal diameters) plus a built-in software. Each needle is connected to a syringe pump through Teflon tubes. One needle (70 μm internal diameter) is attached to a syringe containing FC40 and it is used for jet-printing; this is the jetting needle. The other needle (255 μm internal diameter) is attached to a syringe normally with ethanol to guarantee sterility as this needle is used to handle samples; this is the dispensing needle. Needles can be moved above the surface of a Petri dish as pumps infuse liquids.

In our design, 21 dumbbells were printed in each dish by the jetting needle on d -7. Thereafter, every volume infusion/removal from chamber was operated by the dispensing needle. Every operation on one chamber of dumbbells (either left or right), was repeated to all dumbbells in the dish before starting any task on the opposite chamber.

### Generation of iPSC-derived post-mitotic cortical neurons

This is the yellow pathway in Figure 3. Generation of CNs from two iPSC control lines from healthy human patients (EBiSC409 Cat# STBCi101-A, RRID:CVCL_RD71), and SFC856-03-04 (RRID:CVCL_RC81) was adapted from an established protocol ^26^. In brief, neuronal development was induced with dual SMAD inhibitors 10 μM SB431542 (Tocris) and 118 nM LDN (Sigma) for 25 days (d -33 to d -8). These progenitors (d -8) were transduced with lentiviruses that encoded a doxycylin-inducible tetO promoter driving constitutive expression of rtTA and mouse neurogenin-2 (Ngn2) ^41^. A puromycin resistance gene was also co-expressed to select for cells expressing Ngn2. (See Data Availability for more information about this differentiation protocol).

### Generation of iPSC-derived post-mitotic medium spiny neurons

This is the red pathway in Figure 3. Two iPSC control lines (SFC156-03-01 and SFC856-03-04) were differentiated into MSNs using conditions modified from established protocols ^27^. In brief, neural induction of sub-pallial identity was initiated by dual SMAD inhibitors – 10 μM SB431542 (Tocris) and 118 nM LDN (Sigma) – and inhibition of WNT signalling – 4 μM XAV (Tocris) – from d -16 to d -8. From d - 8 to d 0, SB431542 was removed and neurogenesis in the culture mediated by LDN and XAV. Activin A (25 *µ*g/ml; SKU# SRP3003), the key regulator of TGF-β signalling, was added from d -4 to d 0. (see Data Availability for more information about this differentiation protocol).

### Fabrication of dumbbells

This is the blue pathway in Figure 3 from d -8 to d -6. On d -8, 6 cm Petri dishes were pre-coated with 7 ml of poly-D-lysine (0.01 mg/ml) overnight. On d -7, dishes were washed twice with PBS and subsequently loaded with Neurobasal medium (1 ml; ThermoFisher) supplemented with 1x B27 (ThermoFisher). After at least 5 minutes, medium was manually removed to leave a thin layer (∼50 *µ*l) attached to the bottom of the dish, and FC40 (∼2 ml) gently pipetted on to this thin layer. An array of 7×3 dumbbells was jet-printed using a fluid printer (iotaSciences Ltd.) ^22^. When used with cells, dumbbell chambers were each loaded with 2 *µ*l Geltrex™ (0.46 mg/ml; ThermoFisher) and incubated at 37°C overnight (to d -6).

### Maturation media used for culturing in dumbbells

Cortical maturation medium (CN medium) used from d -6 to d 25 and beyond: Neurobasal, 1X B27 with vitamin A, 1X Glutamax, penicillin/streptomycin (50 U or mg per ml) (1:200), 1 mg/ml doxycycline (1:1000), 10 ng/ml BDNF, 10 ng/ml NT-3 (1:1000), 200 ng/ml laminin (1:5000), 200 mM ascorbic acid (1:1000). This corresponds to Ngn2 base medium from protocol dx.doi.org/10.17504/protocols.io.bp2l69qr5lqe/v1.

Cortical maturation medium + 10x BDNF (BDNF medium) used from d 0 to d 25 and beyond: CN medium with 100 ng/ml BDNF. Striatal maturation medium (MSN medium) used from d 0 to d 8: DMEM/F12 basal medium, 1% MEM Non-Essential Amino Acids (NEAA), 1% L-glutamine, 1x B27 without vitamin A, 1% penicillin/streptomycin (P/S), 0.05% β-mercaptoethanol. This corresponds to striatal maturation medium 1 from protocol dx.doi.org/10.17504/protocols.io.eq2ly79prlx9/v1.

Striatal maturation medium (MSN medium) used from d 8 to d 25: 50% DMEM/F12 basal medium, 50% Neurobasal, 1% MEM Non-Essential Amino Acids (NEAA), 1% L-glutamine, 1x B27 plus vitamin A, 1% penicillin/streptomycin (P/S), 0.05% β-mercaptoethanol. This corresponds to striatal maturation medium 2 from protocol dx.doi.org/10.17504/protocols.io.eq2ly79prlx9/v1.

### Plating CNs in left-hand chambers and transduction

This is the blue pathway from d -6 to d 0. On day -6, 2 *µ*l was removed from each chamber, 4 *µ*l cortical-maturation media deposited in the right-hand chamber (to create a positive pressure gradient toward the left chamber), and post-mitotic CNs plated (1 *µ*l containing 13,000 cells in cortical maturation medium) in each left-hand chamber. On day -3, CNs were transduced by adding 2 *µ*l lentivirus encoded NGN2-GFP in cortical maturation medium (50 *µ*l of viral stock/2 *µ*l) to the left-hand chamber, and 1 *µ*l fresh medium without lentiviruses to the right-hand one (to give ∼4 *µ*l/chamber). On d -2, 4 *µ*l were removed from each chamber and 4 *µ*l fresh medium added back (to wash away free lentivirus).

### Varying conditions on d 0

On d 0, both chambers initially contained ∼4 *µ*l cortical maturation medium, with CNs in the left-hand one. Then, contents of the distal chamber were varied to give the 3 conditions (CNs-CN medium, CNs-BDNF medium, and CNs-MSNs). For all conditions, 4 *µ*l was removed from both chambers, and 4 *µ*l CN medium added to the left-hand chamber. Next, 1 *µ*l of either CN medium, or CN medium plus 100 ng/ml BDNF, or a MSNs suspension (13,000 cells in MSN medium) was added to the distal chamber. As before, a pressure difference confined MSNs and BDNF to right-hand chambers.

### Culturing CNs in dumbbells without MSNs after d 0

This is the blue pathway from d 0 to d 25 and beyond for the CNs-CN medium and CNs-BDNF medium conditions. On d 2, both chambers were completely emptied (removing ∼4 *µ*l from each chamber). Then, 4 *µ*l CN medium was added to the CN chamber and 4 *µ*l of either CN or BDNF medium to the distal one (to give 4 *µ*l/chamber). Every 48 h thereafter, half the specified medium present in a chamber was replenished by removing 2 *µ*l spent medium and then adding back 2 *µ*l fresh medium of the same kind.

### Culturing CNs in dumbbells with MSNs after d 0

This is the blue pathway from d 0 to d 25 and beyond for the CNs-MSNs condition. On d 2, both chambers were completely emptied (removing ∼4 *µ*l from each chamber). Then, 4 *µ*l of fresh CN medium was added to CN chamber and 4 *µ*l of MSN medium containing 200 nM cytosine arabinoside (araC) to arrest proliferation of any non-neuronal cells in the population to the distal one (to give 4 *µ*l/chamber). Every 48 h thereafter, half the medium in each chamber was changed by withdrawing 2 *µ*l and adding an equal volume of fresh medium; this gradually diluted araC over MSNs. On d 8, a full medium change is done to switch MSNs to a new MSN medium (see Maturation media used for culturing in dumbbells).

### Axotomy

Fluid walls were destroyed by gently pouring all FC40 out of the dish followed by two washes with cortical maturation medium (care is taken to prevent cell/axon peeling). Directed axotomy was performed automatically using the fluid printer by modifying an existing procedure ^42^. A 1 ml glass syringe (Hamilton) filled with Neurobasal medium (Thermofisher) was connected via Teflon tubes to the jetting nozzle. Then, a medium jet was ejected (480 μl/min) from the nozzle (70 μm inner diameter) held 0.3 mm above the dish as the traverse moved (960 mm/min) the nozzle in a straight line perpendicular to the axons’ main direction of outgrowth (Figure 4B). Following axotomy, dishes were re-filled with fresh FC40, and new fluid walls jet-printed around the original footprint. The new dumbbells had larger footprints (chamber area = 3.5×3.5 mm^2^, conduit length = 0.5 mm, width ∼400 *µ*m) to avoid damaging attached cells/axons. 4 μl fresh cell medium was deposited into each chamber.

Figure 6 summarises data obtained from 2-3 differentiations per cell line, and measurements from 100-150 dumbbells/differentiation. Another ∼30% dumbbells were discarded due to incomplete severing or clearing of axons in the axotomised area of the conduit, and/or incomplete rebuilding of new dumbbells (which results in media leakage). [Fluid walls are almost always built successfully on virgin Petri dishes, but success rates are lower when building on dishes that have been covered with a thin skim of medium overlaid with FC40 from d -7 to d 20.]

### Imaging

All fluorescent live-cell images of dumbbells were taken with a digital single-lens reflex camera (Nikon D7100 DSLR) connected to an epifluorescence microscope (Olympus IX53). Images were analysed using Cell Profiler 3.8 (RRID: SCR_007358) to describe and quantify axon outgrowth and regrowth.

### Immunostaining

All FC40 was discarded from the dish, cultures washed twice with PBS, fixed (2% paraformaldehyde, 20 mins at room temperature, RT), and washed three times with PBS. Fixed samples were then incubated (80°C, 5 min) in citrate buffer pH 6.0 (ThermoFisher), and left for 10 mins at RT. Permeabilization and blocking were performed concurrently in PBS, 10% donkey serum, and 0.01% Triton X-100 for 10 mins. Following incubation with primary antibody in PBS and 10% donkey serum overnight at 4°C, samples were washed with PBS, and incubated in species-appropriate Alexa Fluor© secondary antibody in PBS with 10% donkey serum for 1 hour at RT. Supplementary Table 1 list antibodies used. Images are acquired on an Invitrogen EVOS ™ FL Auto (ThermoFisher) cell-imaging system, and processed using ImageJ (RRID:SCR_003070) ^43^.

### Measuring pressures and volumes in chambers

Dumbbells were printed in wide rectangular plates (Thermo Scientific™ Nunc™ Rectangular Dishes single well) to improve imaging from the side, as the curved plastic walls of 6 cm dishes distort views. Chambers were jet-printed as before after filling a dish with ∼5 ml DMEM + 10% FBS, removing all but a thin film, and overlaying ∼10 ml FC40. Using a fluid printer modified to host rectangular dishes (Hylewicz CNC-Technik). The same medium was infused into chambers by a syringe pump (PhD Ultra, Harvard Apparatus) equipped with a 50 μl glass syringe (Hamilton) connected to a blunt metal needle (33G blunt NanoFil™ needle, World Precision Instruments) through a Teflon tube (Zeus Company Inc.) to generate the desired initial volume/pressure difference. Volumes were dispensed following the same sequence used during experiments with neurons. Thus, 4 μl were initially infused in the right-hand chamber, followed by 1 μl into the left-hand one. Images of the two chambers equilibrating were recorded from the side every 30 min for 24 h using a camera (FTA1000 B Class, First Ten Angstrom) placed perpendicular to the connecting conduit (Figure 2B). Chambers heights are measured using FTA32 software (First Ten Angstrom, RRID: SCR_024392) and the outer diameter of the dispensing needle (210 μm) as a scale reference. Heights were converted into pressures and volumes (Figure 2D) using equation 1 and equation 2, respectively.

### Statistical analysis

All data were presented as mean ± standard error of the mean (SEM) unless otherwise stated. Raw data were tested for normality and statistical comparison of the means is performed using one-or two-way ANOVA with Bonferroni post-hoc test; a difference is considered significant if p < 0.05. All statistical analyses were performed on GraphPad Prism 6.0 (GraphPad Software, RRID:SCR_002798).

## Data Availability

The data that support the findings of this study are deposited on Zenodo (DOI doi.org/10.5281/zenodo.7924431). All details of the antibodies, cell lines, and software used in this work are available on Zenodo (DOI doi.org/10.5281/zenodo.7924431). Protocols associated with this work can be found on protocols.io (DOI dx.doi.org/10.17504/protocols.io.36wgqjwwxvk5/v1). The custom G-Code scripts used in this study to jet-print the microfluid-walled dumbbells are available at https://github.com/craggASAP/microfluid_axotomy.git.

## Author Contributions

F.N. and Q.D. conceived the project. F.N. and Q.D. designed, performed, and analysed all experimental data. P.R.C, E.W., and R.W.M supervised the study. F.N. and Q.D. prepared the first draft of the manuscript. All authors reviewed the manuscript and approved its submission.

## Acknowledgements

This work was supported by iotaSciences Ltd and the Engineering and Physical Sciences Research Council through EP/R513295/1 (who both provide financial support to F.N.). Q.D. was supported by a National Science Scholarship from Agency for Science, Technology and Research in Singapore. This research was funded in part by Aligning Science Across Parkinson’s [ASAP-020370] through the Michael J. Fox Foundation for Parkinson’s Research (MJFF) and in part by the Monument Trust Discovery Award from Parkinson’s UK (J-1403). The work was supported by a National Institute for Health Research-Medical Research Council Dementias Platform UK Equipment Award (MR/M024962/1) to R.W.M. For the purpose of open access, the author has applied a CC BY public copyright license to all Author Accepted Manuscripts arising from this submission. We thank Nora Bengoa-Vergniory and Cristian Soitu for contribution to early stages of the project and thank Ajantha Abbey for his generous donation of cortical neuron media.

