## Supplementary Information for "A fluid-walled microfluidic platform for human neuron microcircuits and directed axotomy"

**Determination of concentration gradients in dumbbells**

Consider the dumbbell in Supplementary Figure 1, where chambers are filled with equal volumes, and transport of molecules through the conduit is driven only by diffusion to generate a concentration gradient from right to left. Prediction of those gradients represents a crucial step in understanding the roles of target-derived signals like BDNF or other MSN-derived factors. Using Fick’s 2^nd^ law, concentration (c(x,t)) in the conduit can be approximated by the solution to the one-dimensional diffusion equation for a semi-infinite medium with constant concentration at the boundary ^1^

$$c(x,t)=C_{0}erfc\left( \frac{x}{2\sqrt{Dt}} \right)$$

(S1)

with $C_{0}$ being the initial concentration in the right-hand chamber, D the diffusion coefficient, and x and t being space and time respectively. Equation S1 allows estimation of the time necessary for the gradient to reach steady state ($t_{steady}$), arbitrarily quantified as the time required for 30% of $C_{0}$ to reach the other end of the conduit $(x=L), t_{steady}=\frac{1}{D}\left( \frac{L}{2 erfc^{-1}(0.3)} \right)^{2}.$ The value of 30% is chosen so that the maximum error between the gradient profile (Eq. S1) and the linearized approximation is smaller than 10%. Once steady state is reached, the concentration gradient can be considered linear and remain stable for a time $t= t_{linear}$. Deriving from Fick’s 1^st^ law and defining it as the time needed for the concentration in the left chamber to increase by 5% of $C_{0}$, one gets:

$$t_{linear}=1.05\frac{Lm_{mol}}{DA_{c}C_{0}}$$

(S2)

where $m_{mol}$ is the mass of molecules transferred between chambers and $A_{c}$ is the cross-sectional area of the connecting conduit. Unlike conduits with solid walls that have fixed cross sections, ones with fluid walls morph as pressures change. If conduit widths (${2a}_{c}$) remain unchanged (as fluid walls are firmly pinned to the plastic substrate), heights ($h_{c}$) vary depending on pressures applied. In particular, for a cross section like in Supplementary Figure 1B and assuming $h_{c}$<< $a_{c}$ at all times, one can prove:

$$A_{c}=\left( \frac{a_{C}}{2h_{c}} \right)\left[ \left( \frac{a_{C}^{3}}{2h_{c}} \right){sin}^{-1}\left( \frac{2h_{c}}{a_{C}} \right)-a_{c}^{2}+{2h}_{c}^{2} \right]$$

(S3)

where $h_{c}$cannot be considered constant. However, when both chambers enclose the same volume, the dumbbell is in equilibrium and pressure is equal everywhere (Eq. (1)). Such equivalence allows derivation of a relationship between chamber and conduit heights:

$$h_{C}=\frac{(2a_{c}^{2} h_{chamber})}{(a_{chamber}^{2}+h_{chamber}^{2})}- \Delta\rho ga_{c}^{2}h_{chamber}$$

(S4)

With Equation 2, this directly relates chamber volumes and areas of conduit cross sections. Equation S4 has been derived assuming no hydrostatic head of pressure acts in the conduit.  In Table S1, conduit heights and areas are computed for a dumbbell containing 4 μl in each chamber and diffusion parameters are calculated for transport of BDNF ($D=12.6 x {10}^{-7} cm^{2}/s$)^2^. Values are computed for dumbbell geometries used ($m_{mol}=0.02 ng$; growth: $h_{chamber}=1.07 mm, a_{chamber}=1.42 mm$; regrowth: $h_{chamber}=0.85 mm, a_{chamber}=1.67 mm$) pre-axotomy to compare cortical axonal growth, and post-axotomy to analyse regrowth. Supplementary Figure 1C illustrates the linear concentration gradient of BDNF at $t= t_{linear}$, so that its concentration in the right chamber equals $C_{0}=100$ ng/ml and the one in the left chamber is 15 ng/ml (10 ng/ml initially present in the chamber plus 5 ng/ml transferred over $t_{linear}$,).

**Supplementary Table 1. Geometrical parameters of dumbbells with different sizes and respective diffusion times of BDNF**

| **Bio-assay** | **Conduit geometry** | | | | **Diffusion parameters** | |
| --- | --- | --- | --- | --- | --- | --- |
|  | $\mathbf{length [mm]}$ | $\boldsymbol{width [\mu m]}$ | $\mathbf{height}\boldsymbol{[\mu m]}$ | $\boldsymbol{area [\mu}\mathbf{m}^{\mathbf{2}}\mathbf{]}$ | $\mathbf{t}_{\mathbf{steady}} \left[ \mathbf{min} \right]$ | $\mathbf{t}_{\mathbf{linear}}\mathbf{[d]}$ |
| outgrowth | 1 | 200 | 5 | 624 | 60 | 30 |
| regrowth | 0.5 | 400 | 13 | 3513 | 15 | 5 |

**Supplementary table 2**: Primary antibodies

| **Antibody** | **Target** | **Host organism** | **Dilution** | **Source** | **Cat #** | **RRID** |
| --- | --- | --- | --- | --- | --- | --- |
| Anti-DARPP32, monoclonal | DARPP32 | Donkey | 1:250 | Abcam | ab40801 | RRID:AB_731843 |
| Anti-DARPP32, polyclonal | DARPP32 | Donkey | 1:250 | Sigma-Aldrich | HPA048630 | RRID:AB_2680468 |
| Anti-MAP2, Polyclonal | Microtubule-associated protein 2 | Donkey | 1:250 | Abcam | ab92434 | RRID:AB_2138147 |
| Anti-SMI312, monoclonal | Neurofilament marker (pan axonal, cocktail) | Donkey | 1:250 | Biolegend | 837904 | RRID:AB_2566782 |

**Supplementary Table 3**: Secondary antibodies

| **Antibody** | **Target** | **Host organism** | **Dilution** | **Source** | **Cat #** | **RRID** |
| --- | --- | --- | --- | --- | --- | --- |
| Alexa-Fluor 647 | IgG Mouse | Donkey | 1:1000 | Invitrogen | A31571 | RRID:AB_162542 |
| Alexa-Fluor 647 | IgG Rabbit | Donkey | 1:1000 | Invitrogen | A-31570 | RRID:AB_2536180 |
| Alexa Fluor 555 | IgG Chicken | Donkey | 1:1000 | LifeTechnology | A78949 | RRID:AB_2921071 |
| DAPI (4',6-Diamidino-2-Phenylindole, Dilactate) | DAPI | NA | 1:1000 | Thermo Fisher Scientific | D1306 | RRID:AB_2307445 or RRID:AB_2629482 |

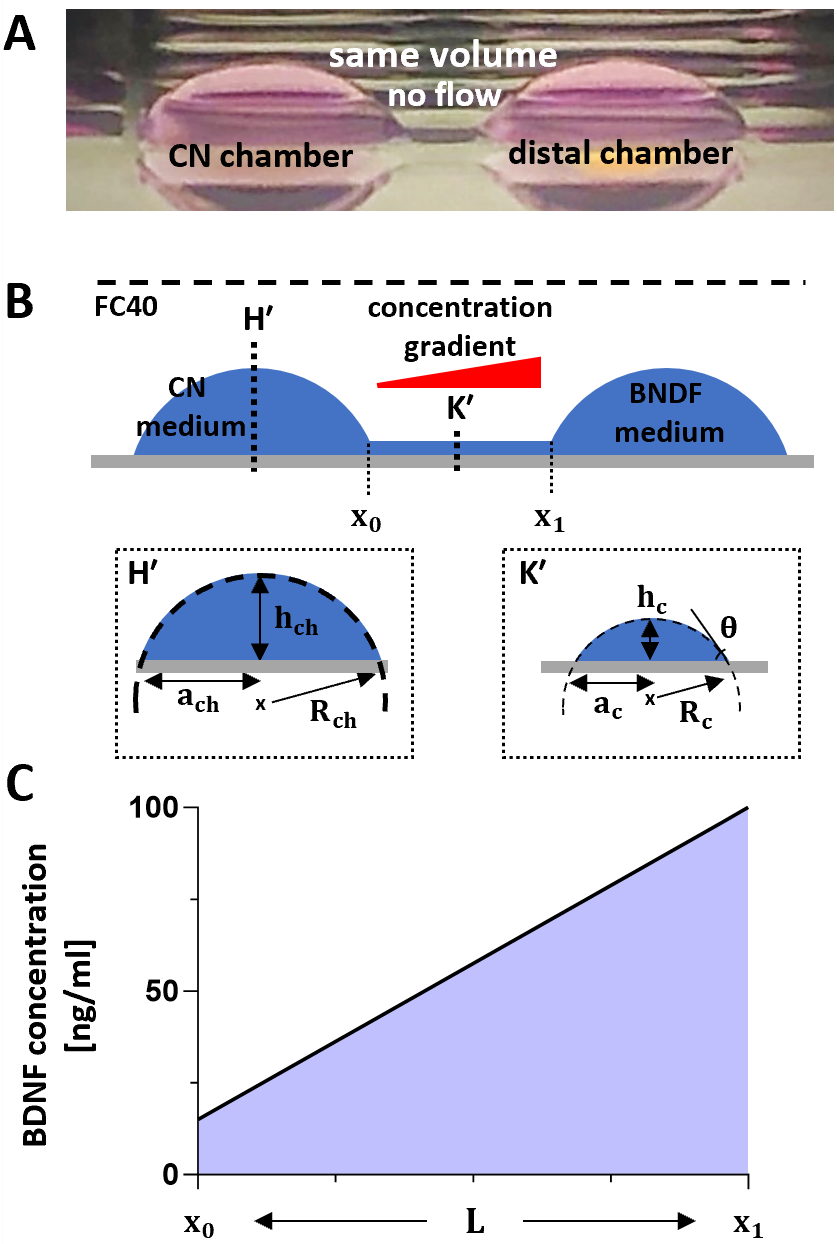

**Supplementary Figure 1. Diffusion gradients in fluid-walled dumbbells.**

**(A)** Side view of dumbbell with 4 μl in each chamber. In all examples shown here and elsewhere, the left-hand one hosts cortical neurons.

**(B)** Schematic of dumbbell with equal volumes. In this case, BDNF molecules diffuse through the conduit, so any axons from CNs will sense a concentration gradient. Lower panels show central cross sections of a chamber and connecting conduit which have shapes of circular segments (as they are bounded by liquid interfaces).

**(C)** Linear approximation of BDNF concentration gradient inside the conduit at $t= t_{linear}$ , when the left chamber ($x<x_{0}$) contains cortical maturation medium (initially $C_{BDNF}$ = 10ng/ml) and the right one ($x>x_{1}$) contains cortical maturation medium with 10-fold BDNF concentration.

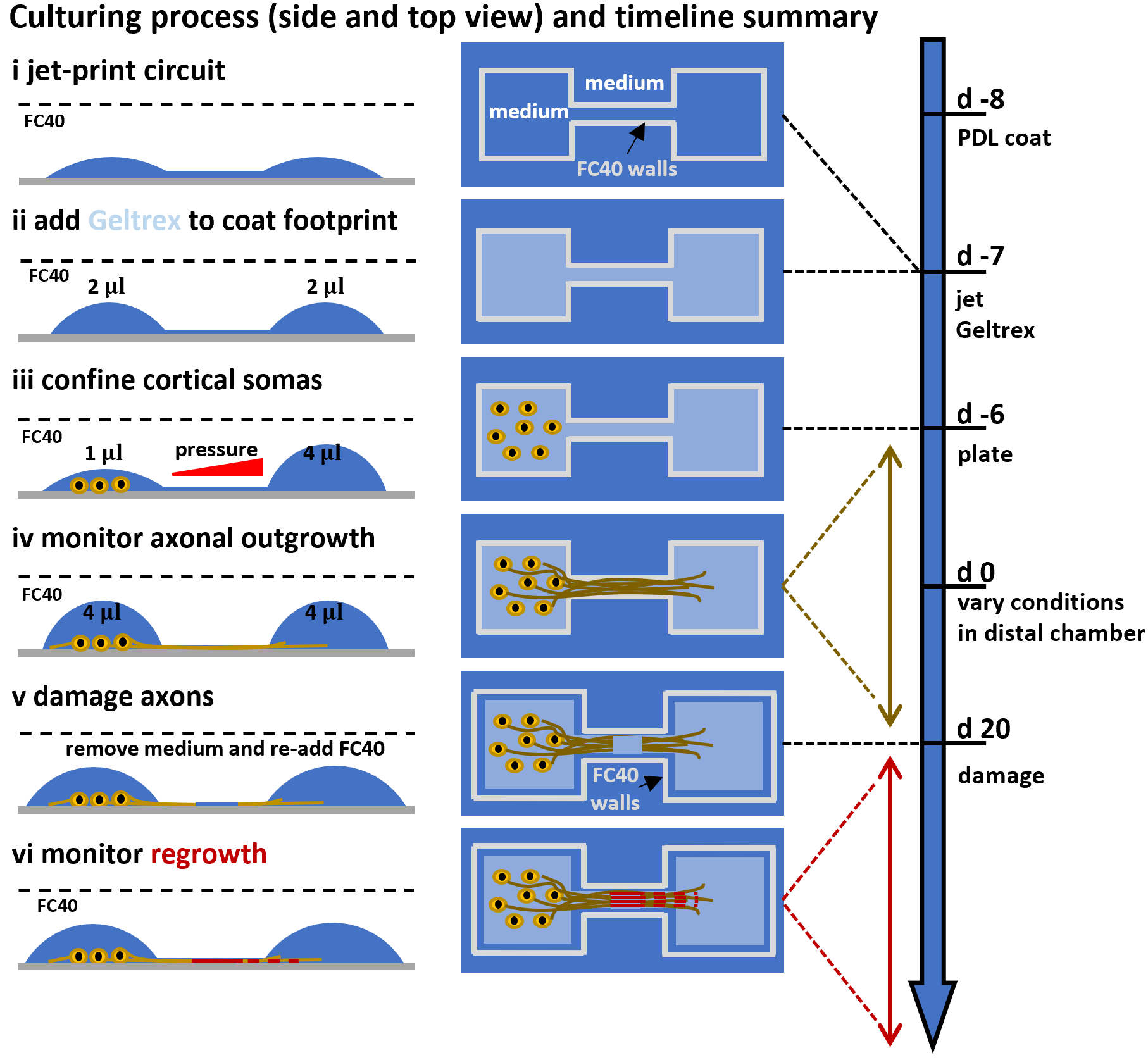

**Supplementary Figure 2. Culturing neurons in fluid-walled dumbbells.**

Overview of most relevant steps followed to culture CNs and perform axotomy in fluid walled dumbbells (top view, side view, and timeline). The culturing protocol is summarised (from d -8 and d 20) between (i) and (iv). The axotomy assay (from d 20 onwards) is shown in (v) and (vi).

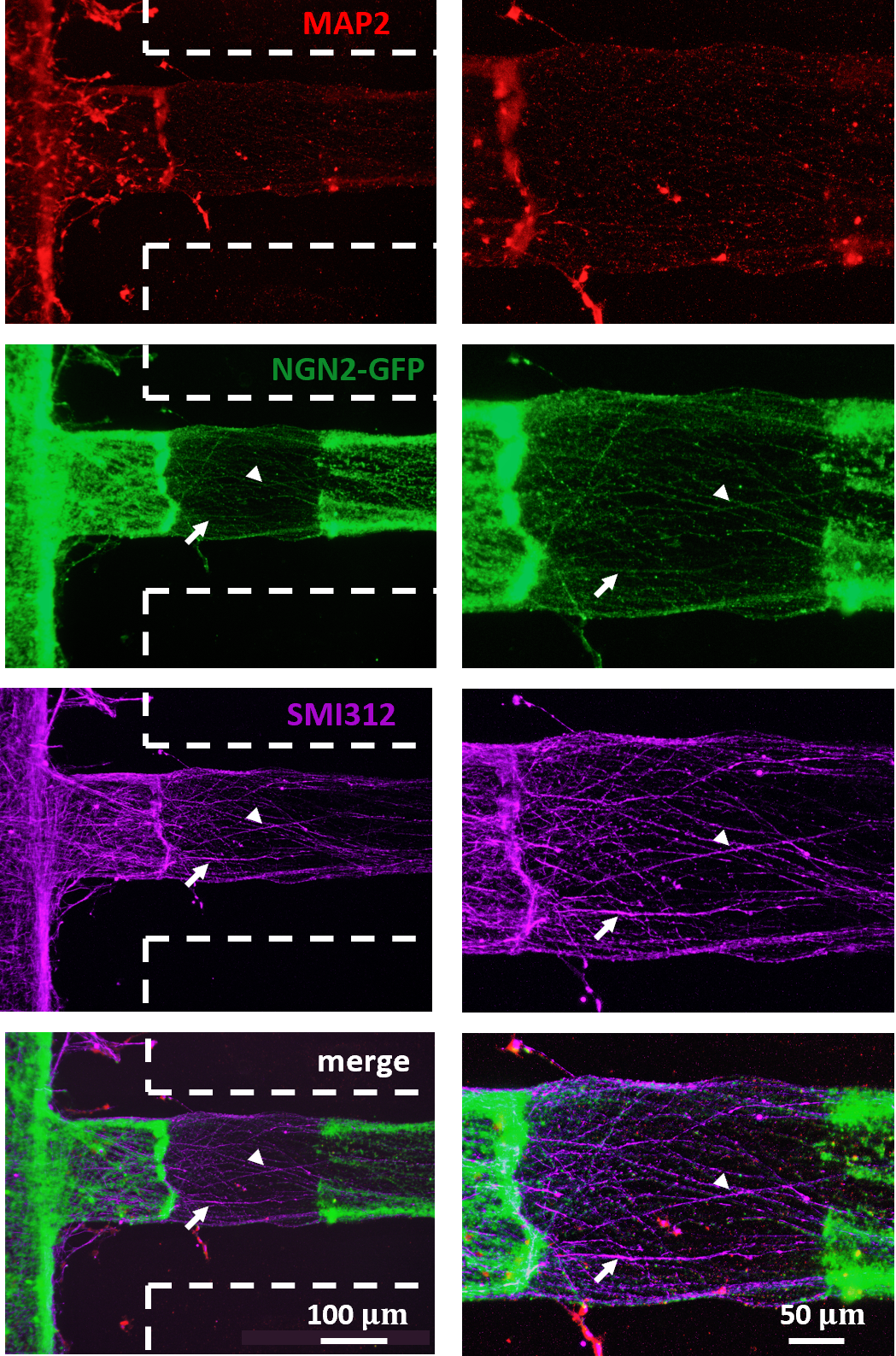

**Supplementary Figure 3. Regeneration after axotomy.**

NGN2-transfected CNs project axons through the conduit. After 26 days of culture, dumbbells are destroyed, axons damaged and new dumbbells re-built to monitor regrowth of axons. Immunostaining images captured 5 days post axotomy show neuronal-domain markers (MAP2 – red – dendrites, SMI312 – purple – axons) and CN marker NGN2-GFP (green). All regenerated neurites are cortical as they express SMI312 and NGN2-GFP, while dendritic MAP2 is absent (arrow and arrowhead mark two examples). Dashed lines indicate the approximate position of the edges of the newly-built footprint; most regenerated neurites are confined to the footprint of the original dumbbell and the Geltrex coat.
